# Wide distributions and cryptic diversity within a *Microstomum* (Platyhelminthes) species complex

**DOI:** 10.1101/290429

**Authors:** Sarah Atherton, Ulf Jondelius

## Abstract

*Microstomum lineare* is a common species of fresh and brackish waters found worldwide. Three genes (18S, CO1, ITS) were sequenced from specimens of *M. lineare* collected from four countries and the levels of cryptic diversity and genetic structuring was assessed. Results showed *M. lineare* has very wide haplotype distributions suggesting higher than expected dispersal capabilities. In addition, three new species were described on the basis of molecular taxonomy: *Microstomum artoisi* sp. nov., *Microstomum tchaikovskyi* sp. nov., and *Microstomum zicklerorum* sp. nov.

## Introduction

Freshwater habitats are thought to have discrete boundaries (Poff 1997; Bohonak & Jenkins 2003), and yet many limnic microinvertebrates have reportedly wide geographical distributions in spite of habitat isolation (Gómez *et al.* 2000; Page & Hughes 2007). Species that maintain broad distributions may be expected to possess high dispersal capabilities, and indeed many species of zooplankton and other freshwater invertebrates share life-history characteristics that would promote rapid dispersal (Havel & Shurin 2004). For instance, a wide variety reproduce asexually, thus allowing a local population to be founded by a single individual only (Bell 1982), or are able to form dormant eggs that provide a long-term egg bank and can resist adverse conditions of overland transport (Mellors 1975; Dodson & Frey 2001; De Meester *et al*. 2002).

However, re-examination of taxa using molecular tools revealed many instances where local populations of freshwater invertebrates are actually genetically divergent (Pfenninger & Schwenk 2007). Many nominal species traditionally seen as cosmopolitan are now recognized as assemblages of morphologically similar or identical cryptic species that are more regionally restricted (reviewed by Bickford *et al.* 2007). It is becoming increasingly evident that species boundaries cannot be determined based on morphology alone; analysis of molecular data is essential.

The utility of DNA sequence data to uncover cryptic lineages and delineate species of micrometazoans has been demonstrated by numerous studies (e.g. Casu & Curini-Galletti 2004; Fontaneto *et al.* 2011; Kieneke *et al*. 2012; Tulchinsky *et al.* 2012; Meyer-Wachsmuth *et al.* 2014) but can be problematic due to its sensitivity to the degree of sampling and the frequent overlap between intra- and interspecific variation (DeSalle *et al.* 2005; Sauer & Hausdorf 2012; Meier *et al.* 2006). Several independent delineation tools have been developed to overcome such problems and connect species discovery with genetic data, including haplotype networks based on statistical parsimony (Templeton *et al.* 1992), Bayesian species delineation (Yang & Rannala 2010; Zhang *et al.* 2013), and maximum likelihood approaches with the General Mixed Yule-Coalescent model (Pons *et al.* 2006). Under ideal circumstances, materials can be collected from populations spread across the entire distribution area of a putative species, thus allowing the distribution of haplotypes to be analyzed through population genetics (Jörger *et al.* 2012).

The use of molecular tools to delimit species is only the first step in the taxonomic process and must be followed by the establishment of formal diagnoses and names in order to be truly useful (Jörger & Schrödl 2013). Failing to do so prevents scientists from including the discovered taxa in future research and otherwise causes confusion through differing numbering systems and lack of proper documentation or vouchers. Unfortunately, there is not one unified method of incorporating DNA sequence data into traditional Linnaean classification, and scientists continue to include DNA sequence information in taxonomic descriptions in a variety of ways (reviewed by Goldstein & DeSalle 2011).

*Microstomum* (Macrostomida: Microstomidae) is a widespread genus of very small, free-living flatworms found in marine, brackish, and freshwaters worldwide. *Microstomum* species are difficult to distinguish through traditional methods, since they have few discrete morphological traits and lack reproductive organs during periods of asexual reproduction. The lack of unambiguous morphological characters has resulted in taxonomic problems and has even caused some scientists to shy away from formally describing new diversity (e.g. Janssen *et al.* 2015). Molecular identification tools could be particularly useful in distinguishing members of this problematic taxon and also make the diversity within *Microstomum* available for metabarcoding-based studies of biodiversity.

*Microstomum lineare* is a cosmopolitan species reported from brackish and fresh waters, from countries in every continent except Antarctica (e.g. Fuhrmann 1894; Graff 1911, 1913; Meixner 1915; Steinböck 1933, 1949; Beklemishev 1950; Ax 1957; Luther 1960; Young 1970; Bauchhenss 1971; Heitkamp 1979; Brusa 2006). As in other *Microstomum* species, it reproduces primarily asexually through fissioning, and only develops reproductive organs and eggs in late summer and fall (Heitkamp 1982). The species can show remarkable variation in the few morphological features used to identify it, including its overall body shape and size, which is known to vary based on food availability and reproduction period (Bauchhenss 1971; Heitkamp 1982); body color, which has been reported as colorless, white, yellow, grey, reddish or brownish (Luther 1960); and even amount of pigmentation in the characteristic red eyespots, which have been shown to vary from large bright red slashes at the frontal end to non-existence depending on light intensity (Bauchhenss 1971). In addition to its purportedly global distribution, the species has a reportedly wide tolerance for differing environmental conditions, such as levels of salinity, temperature, calcium, oxygen, and pH (Kaiser 1969; Young 1973; Karling 1974; Heitkamp 1982). The aims of the present study are to assess for the first time the level of cryptic diversity in *Microstomum lineare* and to examine the degree of genetic structuring of populations throughout Sweden and surrounding areas.

## Methods

### Sampling and Collection

A total of 110 specimens of *Microstomum lineare* were collected over a three-year period from the sediments and aquatic vegetation at eleven locations throughout Sweden, three locations in Finland and one location each from Belgium and the United States (Fig. 1). Collection details, including exact coordinates, abbreviations of sampling area names and specimens collected from each, are listed in Table S1. Samples were transported to the laboratory and allowed to stagnate for 24 to 72 hours before the water was removed into a glass petri dish. The water was then searched with a Nikon SMZ 1500 stereomicroscope, and animals were collected by pipetting and identified using a Leitz LaborLux S compound microscope. Following documentation, individual specimens were fixed in 95% ethanol and transported to Naturhistoriska riksmuseet in Stockholm for DNA extraction and analysis.

**Figure 1:**
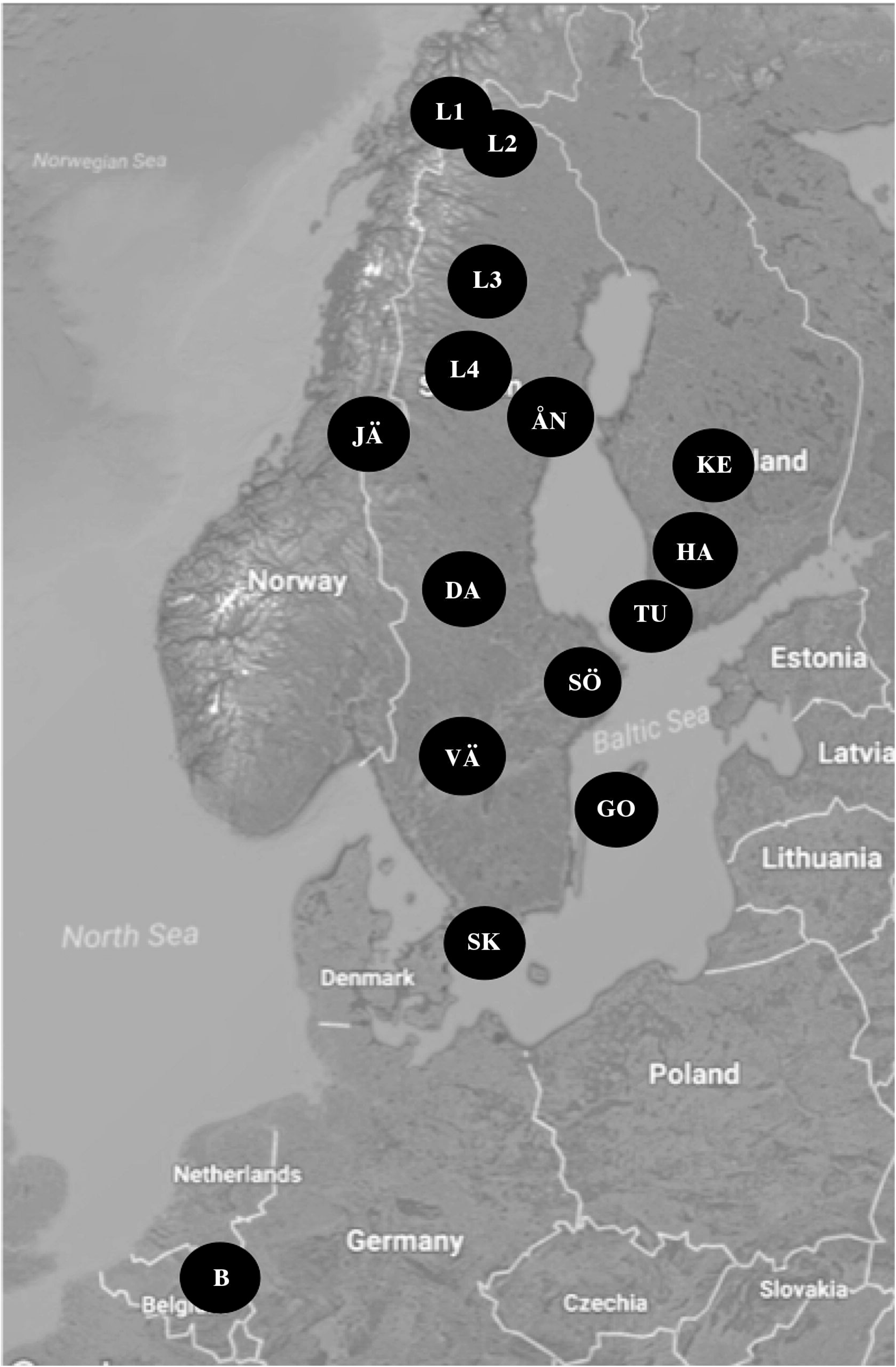
Map of collection locations in Sweden, Finland and Belgium. One additional collection location in the USA (MA, from Massachusetts) was not depicted.

**Table 1:** Number and frequency of haplotypes at each location. Haplotype 6 was found in *Microstomum zicklerorum* sp. n.; haplotype 8 was found in *M. artoisi* sp. n., and haplotype 17-18 were found in *M. tchaikovskyi* sp. n. All other haplotypes stem from *M. lineare*.

|  | DA | JA | ÅN | L4 | L3 | L2 | L1 | B | SK | VÄ | SK | KE | HA | TU | GO | MA | Total | Frequency |
| --- | --- | --- | --- | --- | --- | --- | --- | --- | --- | --- | --- | --- | --- | --- | --- | --- | --- | --- |
| hap1 | 2 | 2 | 3 | 1 | 1 |  |  | 3 | 3 | 3 | 7 | 4 | 1 | 6 | 1 |  | 37 | 0.394 |
| hap2 |  |  |  |  |  |  |  |  |  | 14 |  |  |  |  |  |  | 14 | 0.149 |
| hap3 |  |  |  |  | 6 |  | 7 |  |  |  |  |  |  |  |  |  | 13 | 0.138 |
| hap4 |  |  | 3 |  |  | 5 |  |  |  |  |  |  |  |  |  |  | 8 | 0.085 |
| hap5 |  |  |  |  |  |  |  |  |  | 2 | 2 |  |  |  |  |  | 4 | 0.043 |
| hap6 |  |  |  |  |  |  |  |  |  |  |  |  |  |  |  | 3 | 3 | 0.032 |
| hap7 |  |  |  |  | 2 |  |  |  |  |  |  |  |  |  |  |  | 2 | 0.021 |
| hap8 |  |  |  |  |  |  |  | 2 |  |  |  |  |  |  |  |  | 2 | 0.021 |
| hap9 |  |  |  |  | 1 |  |  |  |  |  |  |  |  |  |  |  | 1 | 0.011 |
| hap10 |  |  |  |  | 1 |  |  |  |  |  |  |  |  |  |  |  | 1 | 0.011 |
| hap11 |  |  | 1 |  |  |  |  |  |  |  |  |  |  |  |  |  | 1 | 0.011 |
| hap12 |  |  |  |  |  |  |  |  |  | 1 |  |  |  |  |  |  | 1 | 0.011 |
| hap13 |  |  | 1 |  |  |  |  |  |  |  |  |  |  |  |  |  | 1 | 0.011 |
| hap14 |  |  |  |  |  |  |  |  |  | 1 |  |  |  |  |  |  | 1 | 0.011 |
| hap15 |  |  |  |  |  |  |  |  |  |  |  |  |  |  | 1 |  | 1 | 0.011 |
| hap16 |  |  | 1 |  |  |  |  |  |  |  |  |  |  |  |  |  | 1 | 0.011 |
| hap17 |  |  |  |  |  |  |  |  |  |  | 1 |  |  |  |  |  | 1 | 0.011 |
| hap18 |  |  |  |  |  |  |  |  |  |  |  | 1 |  |  |  |  | 1 | 0.011 |
| hap19 |  |  | 1 |  |  |  |  |  |  |  |  |  |  |  |  |  | 1 | 0.011 |
| Total | 2 | 2 | 10 | 1 | 11 | 5 | 7 | 5 | 3 | 21 | 10 | 5 | 1 | 6 | 2 | 3 | 94 |  |
| Frequency | 0.021 | 0.021 | 0.106 | 0.011 | 0.117 | 0.053 | 0.074 | 0.053 | 0.032 | 0.223 | 0.106 | 0.053 | 0.011 | 0.064 | 0.021 | 0.032 |  |  |

### DNA extraction, amplification and sequencing

Complete sequences of nuclear 18S and internal transcribed spacer (ITS-1, 5.8S, ITS-2) rDNA as well as a partial fragment of the mitochondrial cytochrome oxidase c subunit 1 (CO1) corresponding to the Folmer region were selected for DNA analyses. The18S ribosomal gene is the most common molecular marker for macrostomorph flatworms on GenBank with 104 entries as of 1 December 2017 and is ideal for comparing diversity at higher taxonomic scales (Giribet 2002), although its low variability in closely related species means that 18S rDNA may underestimate species diversity in some cases (Larsson *et al.* 2008; Tang *et al*. 2012). The ITS and CO1 loci are highly variable and commonly used to identify species and assess intraspecific diversity in Platyhelminthes (e.g. Larsson *et al*. 2008; Lázaro *et al*. 2009; Telford *et al*. 2000; Vanhove *et al*. 2013; Álvarez-Presas *et al*. 2015) and other meiofauna (e.g. Bode *et al*. 2010; Birky *et al.* 2011; Ansian & Tedersoo 2015).

DNA was extracted from whole animals using the DNEasy Blood & Tissue Kit (Qiagen, Valencia, CA) following the manufacturer’s instructions. PCR amplification was performed using 0.2 ml PuReTaq Ready-To-Go PCR beads (GE Healthcare) with 5 pmol each forward and reverse primer and 2 µl DNA. All primers and programs used for amplification and sequencing are listed in Table S2. Products were viewed on a 1% agarose gel, purified using ExoSAP-IT enzymes (Exonuclease and Shrimp Alkaline Phosphatase, GE Healthcare) and sent to Macrogen (Macrogen Europe, Netherlands) for commercial sequencing.

**Table 2:** A hierarchical analysis of molecular variance (AMOVA). Specimens were separated into five different groupings: (1) by country; (2) specimens in Europe vs. USA; and (3) by lineages according to the CO1 ML tree (species of the *M. lineare* complex). d.f. degrees of freedom; ∑ Squares sum of squares; Var Comp variance components; Var % percent of variance; Fix Index fixation index. *p<0.05; **p<0.001

| Source | d.f. | ΣSquares | Var Comp | Var % | Fix Index |
| --- | --- | --- | --- | --- | --- |
| Analysis 1: By Country |  |  |  |  |  |
| Among groups | 3 | 263.176 | V<sub>a</sub> 7.120 | 63.45 | F<sub>SC</sub> 0.105** |
| Among populations in groups | 1 | 19.521 | V<sub>b</sub> 0.429 | 3.82 | F<sub>ST</sub> 0.673** |
| Within populations | 89 | 326.867 | V<sub>c</sub> 3.673 | 32.73 | F<sub>CT</sub> 0.635 |
| Analysis 2: Europe versus USA |  |  |  |  |  |
| Among groups | 1 | 239.355 | V<sub>a</sub> 40.077 | 90.10 | F<sub>SC</sub> 0.126** |
| Among populations in groups | 2 | 23.821 | V<sub>b</sub> 0.557 | 1.25 | F<sub>ST</sub> 0.913 |
| Within populations | 90 | 346.388 | V<sub>c</sub> 3.849 | 8.65 | F<sub>CT</sub> 0.901 |
| Analysis 3: By lineage |  |  |  |  |  |
| Among groups | 3 | 458.949 | V<sub>a</sub> 34.298 | 95.80 | F<sub>SC</sub> -0.117** |
| Among populations in groups | 1 | 1.5 | V<sub>b</sub> -0.176 | -0.49 | F<sub>ST</sub> 0.953 |
| Within populations | 89 | 149.115 | V<sub>c</sub> 1.676 | 4.68 | F<sub>CT</sub> 0.958 |

### DNA alignment and taxonomy

GenBank accession numbers for sequences from all specimens included in the analyses are listed in Table S1. The final datasets included 105 18S (87 *M. lineare*), 121 CO1 (106 *M. lineare*) and 98 ITS sequences (94 *M. lineare*), while the final alignment of the combined dataset included 85 (81 *M. lineare*) sequences. Based on the phylogenetic hypothesis in Janssen *et al*. (2015), sequences of *Myozonaria bistylifera*, *Myozonaria fissipara* and an otherwise unidentified Myozonariinae were downloaded from GenBank and used as outgroups for the 18S and CO1 sequences. As no ITS sequences were available on GenBank for appropriately closely related species at the time of analysis, DNA of two specimens each of *Microstomum rubromaculatum* and *Microstomum hamatum* was sequenced for all three loci and used as the outgroups for the ITS and three-gene concatenated analyses.

Sequence assembly and alignment was performed in MEGA v. 7.0.21 (Kumar *et al*. 2016). Trace files were manually edited, and nucleotide sequences trimmed to equal length. Alignments for each marker were generated using MUSCLE (Edgar 2004) with the default settings. For CO1, sequences were translated to amino acids using the flatworm mitochondrial genetic code (Telford *et al.* 2000), aligned and manually checked for stop codons and reading frame shifts and back translated to nucleotides.

Maximum Likelihood (ML) analysis was performed on all three markers (18S, CO1 and ITS) individually as well as with the three (18S+CO1+ITS) marker concatenated dataset. The best models of sequence evolution were determined using jModelTest2 (Tamura *et al.* 2011) with 3 substitution schemes based upon the AIC criterion (Akaike 1974). The general time reversible model with gamma distribution and proportion invariant sites (GTR+I+G) was selected for the 18S and concatenated datasets, while the general time reversible model with gamma distribution (GTR+G) was selected for CO1 and ITS analyses. All ML analyses were performed with 1000 bootstrap replicates in RaxmlGUI v. 1.5b1 (Silvestro & Michalak 2012).

Species were inferred using the Bayesian implementation of the Poisson Tree Processor (bPTP) in a web-based interface (http://species.h-its.org/ptp) using the default parameters (Zhang *et al*. 2013). bPTP is a tree-based approach to species delimitation that uses number of substitutions in a given tree to model speciation by assuming that the mean number of substitutions per site between species is higher than the number of intraspecific substitutions. The two classes of substitutions are fitted to the tree using a poisson process. bPTP was run multiple times using input trees generated from the ML analyses of the individual CO1 and ITS sequences and from the three-gene concatenated dataset. Additionally, to ensure that results were not influenced by an imbalance of the number of specimens in any given clade, all tests were re-run up to three times with the input tree pruned so that each clade included a maximum of five randomly selected sequences. As recommended, outgroups were not included in the analysis.

### Genetic Diversity

For ITS sequences, a median-joining (MJ) haplotype network (Bandelt *et al*. 1999) was generated with Network 5.001 (www.fluxus-engineering.com). For different groupings of *M. lineare* specimens, number of polymorphic sites (S), number of parsimony informative sites (p), number of haplotypes (h), haplotype diversity (Hd), and nucleotide diversity (π) were calculated in DnaSP (Librado & Rozas 2009), while Tajima’s D were calculated to examine deviations from neutrality. Genetic distances within and among groupings were estimated using the Kimura-two-parameter (K2P) model (Kimura 1980) in Mega 7.02.

Arlequin 3.5222 (Excoffier &Lischer 2010) was used to estimate the degree of differentiation between geographic locations using the pairwise fixation index (F_st_). Isolation by distance (IBD) was tested for specimens collected from Europe using a Mantel test with 1000 iterations. The correlation between pairwise genetic distance (F_st_) and geographic distances was determined with geographic distances approximated as the shortest straight-line distance between sites in km (Table S3).

Finally, a hierarchical analysis of molecular variance (AMOVA) was performed in Arlequin 3.5222 (Excoffier & Lischer 2010) and used to evaluate hypothesized patterns of spatial genetic structure. Specimens were separated into three different groupings: (1) by country; (2) specimens in Europe vs. USA; and (3) by lineage according to ML analyses (Fig. 2).

**Figure 2:**
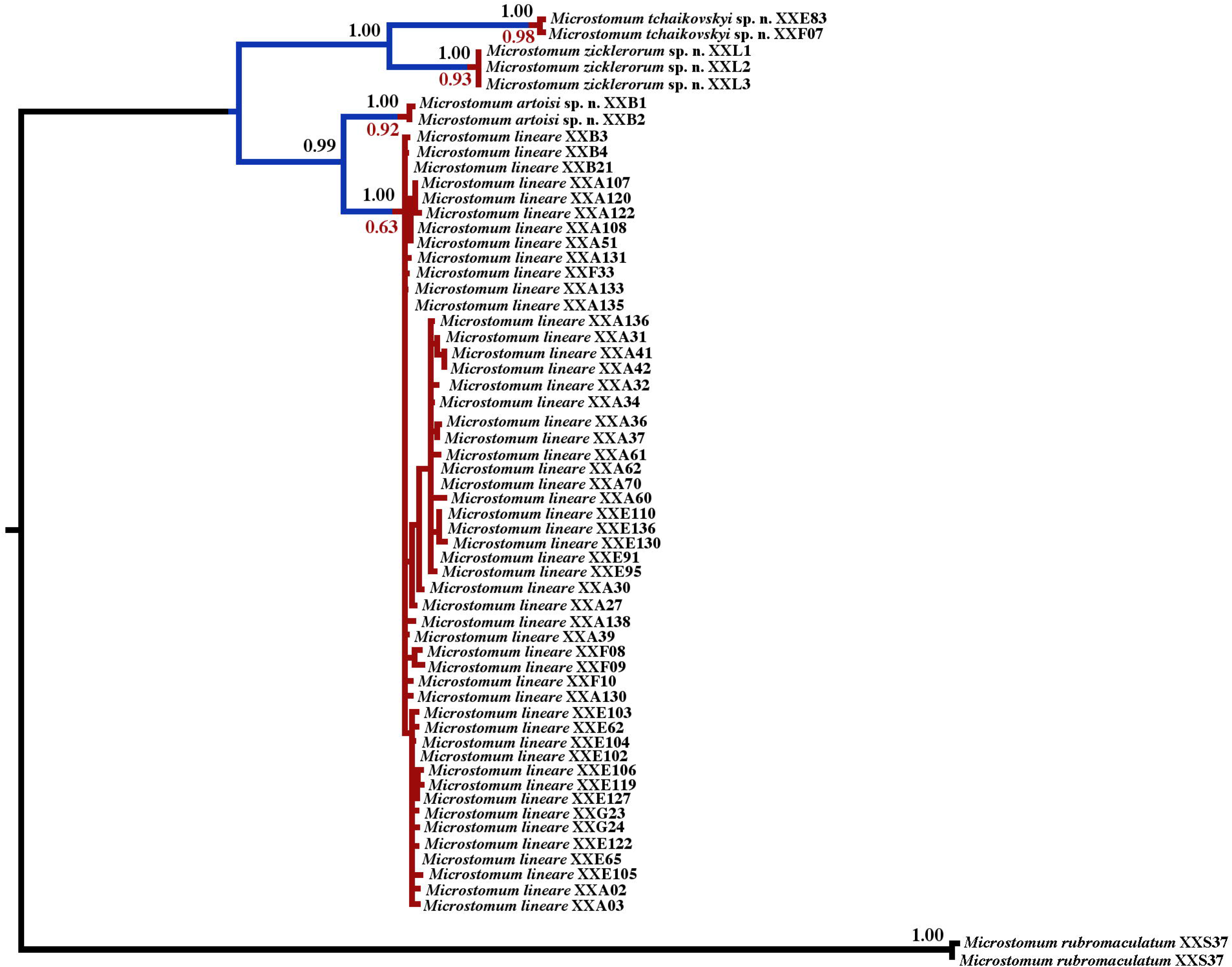
Concatenated 18S, CO1 and ITS1-5.8S-ITS2 gene tree. Specimens are listed with their GenBank accession number. At each node, bootstrap values (black above) and bPTP (red below) values are given. Bars are colored consistent with the results of the bPTP analysis, where the transitions from blue to red represent speciation.

## Results

### Alignments and DNA taxonomy

Alignment of the ITS sequences resulted in a 1064 bp long matrix. The lengths of ITS1, 5.8S and ITS2 ranged from 539-552 bp, 158 bp and 339-341 bp, respectively, and required the insertion of 11 gaps (1-2 bp long) in ITS1 and 2 gaps (1 bp long each) in ITS2. The final aligned sequence matrix included 105 variable sites (10.3%), of which 97 were parsimoniously informative.

The alignment of the CO1 sequences was 654 bp long and required the insertion of three gaps (3-12 bp long) to align the *Microstomum* specimens with the *Myozonaria* outgroups. Alignments of the 18S and three-gene concatenated datasets were 1608 and 3464 characters long, respectively.

Results of the ML analyses of the combined and individual genes were in general accordance. However, as expected, the ITS gene tree showed much better intra-specific resolution than either 18S or CO1. Figure 2 shows the results of the concatenated three-gene phylogeny, while all other trees are provided in Figs. S1-S3.

In all individual and combined ML analyses, specimens of the *Microstomum lineare* complex grouped together to form a monophyletic clade, and within the clade, three specimens from Massachusetts, USA formed a group with moderate (18S gene tree) or very high (ITS, CO1, combined) support (Figs. 2, S1-3). In all analyses except the 18S gene phylogeny, this clade was sister to a second group that included one specimen each from Stockholm, Sweden and Mustalahti, Finland. Additionally, in the ITS gene tree and combined phylogeny, two specimens from Belgium separated to a third clade with very high support, which was sister to the fourth and final clade within the *M. lineare* complex that included specimens from across Sweden, Finland and Belgium.

bPTP results were consistent across all analyses. In the individual and combined datasets with and without taxon pruning*, M. lineare* was divided into four species consistent with the four clades of the ML analyses with moderate (0.63) to high (0.98) support (Fig. 2).

### Genetic Diversity

A total of 19 haplotypes were found for the *M. lineare* complex ITS sequences: 1 in each of 9 locations (DA, JA, L4, L2, L1, MA, SK, HA, TU) 2 in each of 3 locations (B, KE, GO); 3 in 1 location (SÖ); 5 in 2 locations (L3, VÄ); and 6 in 1 location (ÅN). The dominant haplotype (Hap1) was shared by 37 of 94 (39.4%) specimens and occurred in 13 of 16 (81.3%) locations, while each of 11 haplotypes were specific to a single specimen, and 15 haplotypes to a single location. The remaining 7 haplotypes were shared by 2 to 14 (2.1-14.9%) specimens and 2 (12.5%) locations (Table 1).

Pairwise comparisons of different collection locations using the Kimura 2-parameter model of sequence evolution found that ITS sequence divergence ranged from 0 to 0.087 (Table S4). The maximum of 8.7% divergence (88 character changes) occurred between a specimen from Mustalahati, Finland (KE) and Mariestadssjön, Sweden (VÄ), followed by a 7.8% divergence (78 character changes) between two specimens from Massachusetts, USA (MA) and three specimens collected from Norr-Svergoträsket, Sweden (L3). Genetic distances ranged from 0 to 0.037 within locations and from 0 to 0.086 between locations, with the overall highest variations found within KE and between L1 and MA (Table S4).

Figure 3 depicts the ITS *M. lineare* complex haplotype network. The torso includes 15 haplotypes arranged in two star-like configurations each with a main core haplotype (hap1 and hap3). The two central haplotypes differ from each other by 8 mutational steps and from the other haplotypes in the cluster by 1-4 mutational steps. The central haplotypes encompassed 37 of 97 (38.1%) and 13 of 97 (13.4%) of the total specimens. There were 11 (57.9%) unique singleton haplotypes with the highest originating from ÅN (n = 4). Consistent with the ML analyses, Hap 6 (n = 3; collected from MA), Hap 8 (n = 2; collected from B), Hap 17 (n = 1; collected from SÖ) and Hap 18 (n = 1; collected from KE) were distant from the inferred ancestral haplotypes (Hap1).

**Figure 3:**
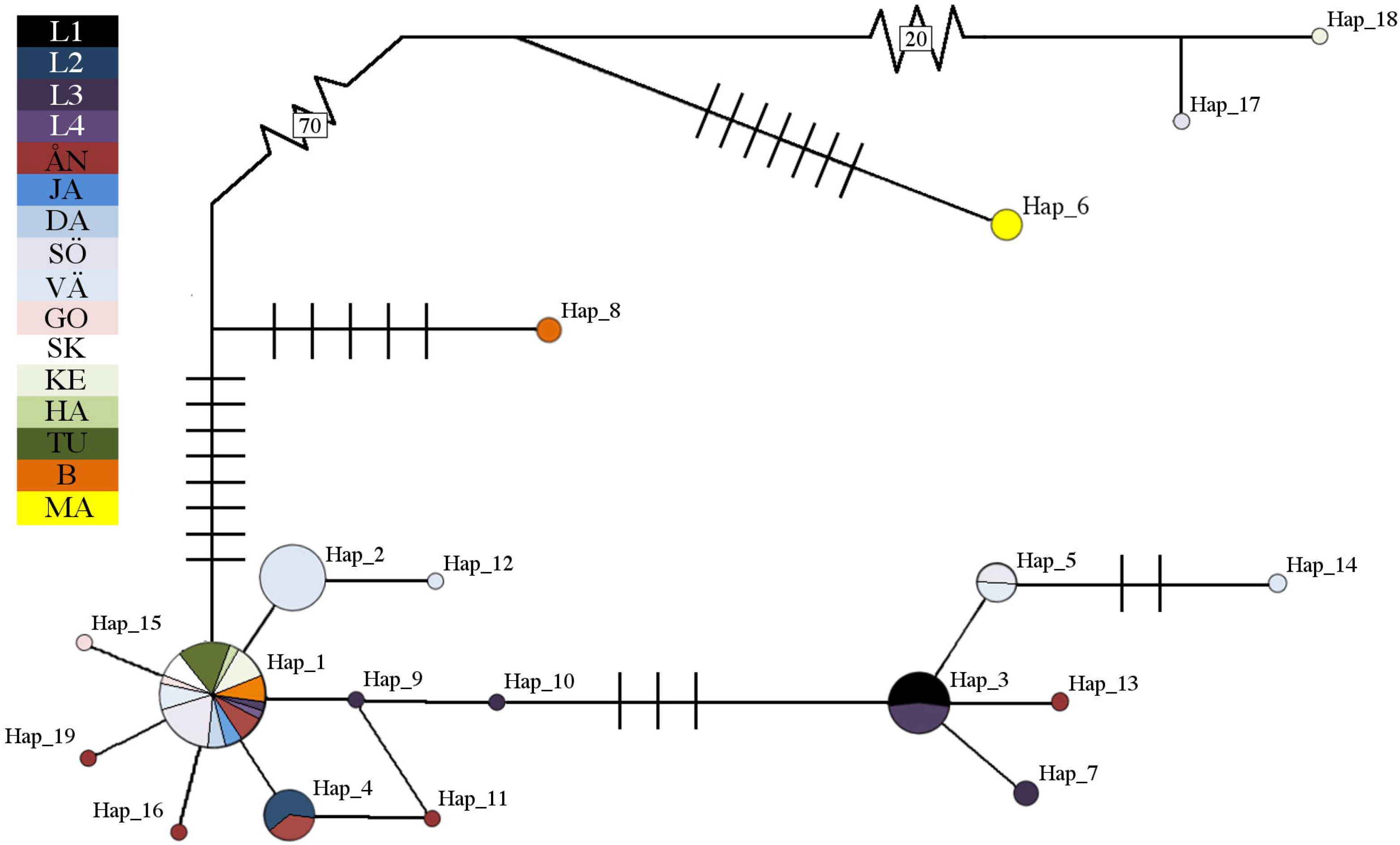
Network Haplotype Map for ITS1-5.8S-ITS2 sequences. Circle size is relative to haplotype frequency. Slashes across haplotype connections indicate additional mutational steps, with large numbers of additional mutational steps indicated by boxed numbers. Key is relative to collection areas as defined in Fig. 1 and Table S1. Hap haplotype

The AMOVA tests found that most of the genetic variation (95.8%) could be explained by variation between inferred ML lineages (Table 2). When specimens from the USA (MA) were separated from the remaining European specimens, between-group variation accounted for 90.1% of the genetic variation, while when the specimens were separated by country, between-group variation accounted for only 63.5%. There was no evidence of isolation by distance (IBD) among European samples (Mantel test; R=0.098 P=0.252).

Table 3 lists the diversity and neutrality indices for ITS sequences of the different groupings of the *M. lineare* complex. Overall, the group inclusive of all studied specimens had a haplotype diversity (hd) index of 0.800 and a nucleotide diversity (π) of 0.01076. When separated by lineage following the ML and network analyses, the largest clade (90 specimens) show hd and π values of 0.767 and 0.00286, respectively. Tajima’s D showed significantly negative results, indicative of population expansion or bottleneck, when all specimens were included in the analysis as well as in the Sweden, Finland, Sweden+Finland and Europe subgroupings.

**Table 3:** Statistics for collected specimens of the *Microstomum lineare* complex. Number of polymorphic sites (S), number of parsimony informative sites (P), number of haplotypes (H), haplotype diversity (Hd) with standard deviation (SD), nucleotide diversity (π), and Tajima’s D (Tajima 1989) were calculated for different population groupings. *M. lineare* does not include the other species of the *M. lineare* complex. *p<0.05; **p<0.001

| Population | No. of Isolates | S | P | H | Hd±SD | $\pi$ | Tajima's D |
| --- | --- | --- | --- | --- | --- | --- | --- |
| All | 94 | 992 | 126 | 19 | 0.800±0.033 | 0.011 | -1.505* |
| Belgium | 5 | 1012 | 13 | 2 | 0.600±0.175 | 0.008 | 1.816 |
| Finland | 12 | 998 | 82 | 2 | 0.167±0.134 | 0.014 | -2.307** |
| Sweden | 74 | 994 | 94 | 16 | 0.831±0.025 | 0.005 | -2.348** |
| USA | 3 | 1004 | 0 | 1 | 0 | 0 | - |
| Europe | 91 | 992 | 98 | 18 | 0.787±0.034 | 0.007 | -2.070* |
| *M. lineare* | 87 | 1008 | 15 | 15 | 0.767±0.036 | 0.003 | -0.283 |

### Taxonomy

Order Macrostomida Karling, 1940

Family Microstomidae Luther, 1907

Genus *Microstomum* Schmidt, 1848

### *Microstomum lineare* (Müller, 1773) Ørsted 1843 (Figs. 4A-C, S4-6)

*Fasciola linearis* Müller, 1773: 67

*Planaria linearis* Müller, 1776: 223

*Derostoma lineare* Duges, 1828: 141

*Anotocelis linearis* Diesing, 1862: 237

*Microstomum caudatum* Leidy, 1852: 350

*Microstomum inerme* Zacharias, 1894: 83

### Materials examined

13 specimens from Finland, 3 specimens from Belgium, 87 specimens from Sweden. Table S1 lists exact dates, times and coordinates. (SMNH XXX; Genbank accession XXX)

### Molecular Notes

COI gene region with reference to Genbank Accession XXXX: 3-A, 57-G; 63-A; 99-T; 258-G; 414-A; 549-A

ITS1, 5.8S, ITS-2 gene region with reference to Genbank Accession XXA02: 29-A; 120-G; 355-T; 819-C; 858-T; 878-T

### Morphological Notes

Body length to approximately 5 mm, maximum 8 zooids. Body color grey, yellowish or clear and reflective of intestines. Anterior end conical, posterior end tail-like with adhesive papillae. Sensory bristles present at extremities. Ciliary pits present.

Two red stripes present in some individuals at dorso-anterior end. Amount of pigmentation variable, from two bright bands to absent (Fig. S4). Some individuals may have two additional faint red pigment stripes ventro-anteriorly (Fig. S4D). Red eye pigmentation may also be present dorsolaterally at the anterior margins of well-developed zooids (Fig. S4F).

Mouth, pharynx, intestine as typical for the genus. Preoral gut present to level of brain. Intestine often with animal food, notably nematodes, cladocerans, ostracods and chironomid larvae (Fig. S6).

Nematocysts present throughout the body.

Reproductive system includes bilateral testes and a single ovary (Fig. S5). Vesicula seminalis circular to eliptical, to approximately 155 µm in diameter, with prostate glands. Stylet spiraled, maximum 125 µm long. Gonopores separate

### Microstomum tchaikovskyi sp. n. (Fig. 4D)

#### Holotype

Vegetative, SWEDEN, Sollentuna, 59°26’26’’N 18°00’03’’E, XX cm depth, 15 September 2016, Atherton, (SMNH-Type-XXXX; Genbank accession XXXX),

**Figure 4:**
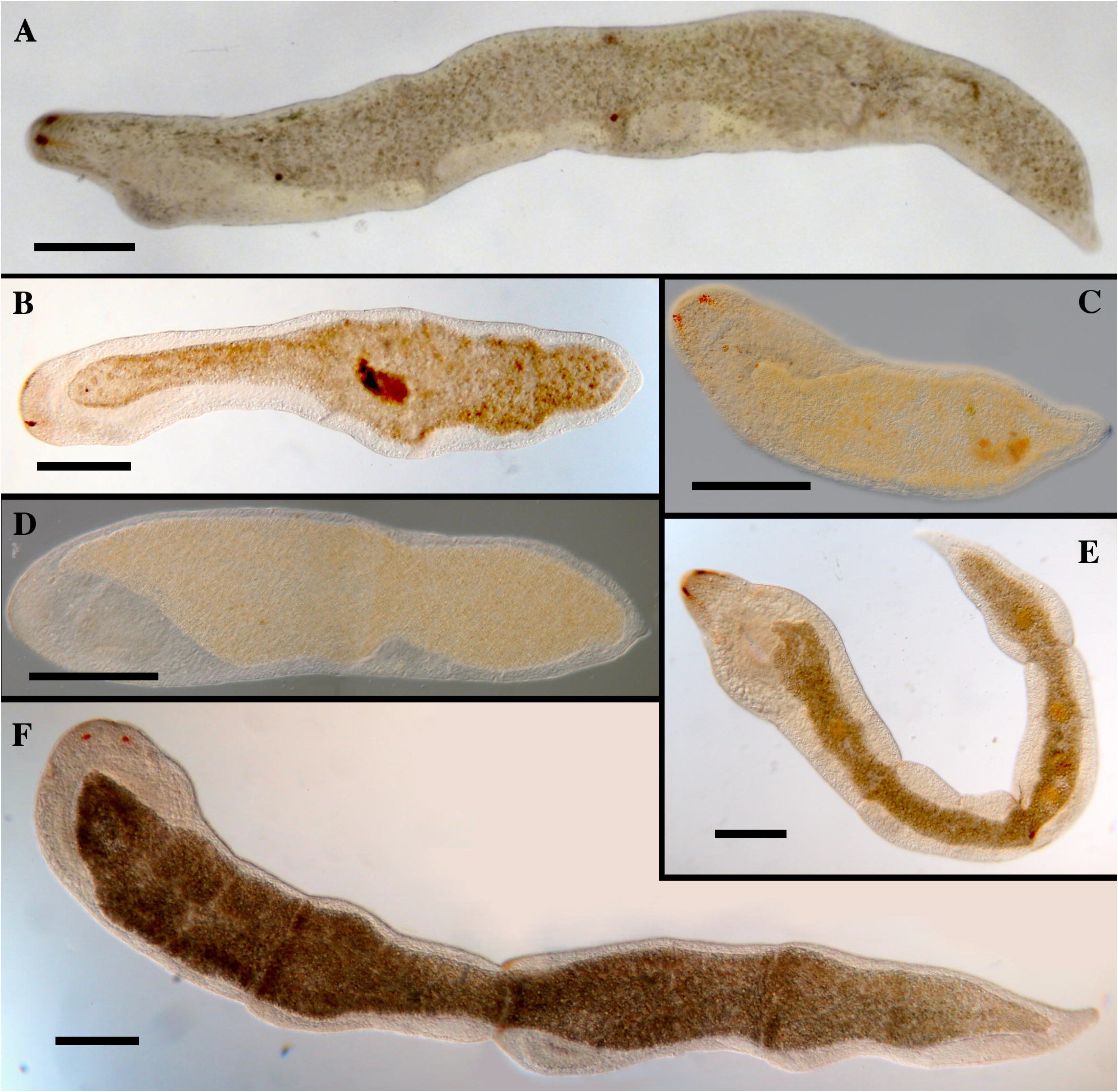
*Microstomum lineare* complex. A. *M. artoisi* sp. n. (GenBank Accession XXXB02) collected from Hoge Kempen Park in Belgium. B. *M. tchaikovskyi* sp. n. (GenBank Accession XXXF07) collected from Tampere, Finland. C. *M. lineare* (GenBank Accession XXXG24) collected from Gotland, Sweden. D. *M. zicklerorum* sp. n. (GenBank Accession XXXL03) collected from Massachusetts, USA. E. *M. lineare* (GenBank Accession XXXF08) collected from Tampere, Finland. F. *M. lineare* (GenBank Accession XXXA116) collected from near Umeå, Sweden. Bar indicated 300 µm for all images.

#### Other material

1 Vegetative, FINLAND, Tampere, 62°01’31’’N 24°41’03’’E, XX cm depth, 5 June 2017(SMNH-XXXX; Genbank accession XXX)

#### Etymology

This species is named in honor of Adrian Tchaikovsky, author, and Pyotr Ilyich Tchaikovsky, composer, whose audiobook and music respectively were much appreciated by the first author while sampling.

### Molecular Diagnosis

COI gene region with reference to Genbank Accession XXXX: 6-T; 15-A; T-16; 18-A; 30-G; 54-A; 69-A; 84-A; 180-G; 186-G; 228-A; 246-C; 279-A; 310-C; 312-T; 318-A; 321-A; 333-A; 351-A; 435-T; 486-G; 540-A; 603-A; 609-1; 625-C; 627-T ITS1, 5.8S,

ITS-2 gene region with reference to Genbank Accession XXA02: 121-T; 218-A; 288-T; 353-G; 358-G; 394-G; 497-T; 533-A; 730-G; 759-C; 768-C; 787-A; 800-T; 845-T; 887-T; 999-T

### Morphological Description

Body length approximately 4 mm, 6 zooids. Body clear with darker intestines. Anterior end abruptly tapering, posterior end tail-like with adhesive papillae. Sensory bristles present. Ciliary pits present at the level of anterior pharynx.

Red eye pigmentation in two stripes, located dorsally at anterior end. May be weak in some individuals.

Preoral gut extending just to level of brain. Mouth, pharynx and intestine as normal for genus. Nematocysts few, scattered across the body.

Reproductive system not seen.

### Microstomum zicklerorum sp. n. (Fig. 4E)

#### Holotype

Vegetative, USA, Massachusetts, North Chelmsford 42°38’13’’N 71°23’30’’W, 25 cm depth, 11 April 2017, Atherton, (SMNH-Type-XXXX; Genbank accession XXXX)

#### Other material

2 Vegetative, Same data as for holotype (SMNH-XXXX; Genbank accession XXX)

#### Etymology

This species is named in honor of Dr. Stephan Zickler, Shannon Zickler, Sophie Zickler, Sabrina Zickler and Stella Zickler.

### Molecular Diagnosis

COI gene region with reference to Genbank Accession XXXX: 24-A; 30-T; 123-T; 165-A; 186-A; 192-G; 201-G; 267-C; 300-C; 318-G; 324-G; 330-A; 336-C; 387-C; 501-A; 525-G; 555-G; 558-A; 588-A; 609-G; 630-C

ITS1, 5.8S, ITS-2 gene region with reference to Genbank Accession XXA02: 197-A; 317-A; 507-T; 829-A; 879-A; 888-C; 1000-A

### Morphological Description

Body length approximately 1.5 mm, 2 zooids. Body clear and reflective of intestines. Anterior end conical, posterior end tail-like with adhesive papillae.

Sensory bristles present. Ciliary pits present with circular openings lateral at the level of mid-pharynx.

Red eye pigmentation extremely weak or not present.

Preoral gut present and extending to level of brain. Mouth and pharynx large leading to strait intestine. Intestine containing desmid algae. Nematocysts present throughout the body.

Reproductive system not seen.

### Microstomum artoisi sp. n. (Fig. 4F)

#### Holotype

Vegetative, BELGIUM, Hoge Kemper Park, 51°00’03’’N 5°40’44’’E, 10 cm depth, 22 June 2017, Atherton & Artois, (SMNH-Type-XXXX; Genbank accession XXXX),

#### Other material

1 Vegetative, BELGIUM, 50°57’45’’N 5°23’22’’E, 5 cm depth, 20 June 2017, Atherton (SMNH-XXXX; Genbank accession XXX)

#### Etymology

This species is named in honor of Dr. Tom Artois.

### Molecular Diagnosis

COI gene region with reference to Genbank Accession XXXX: 39-A; 42-G; 57-A; 81-C; 114-A; 180-T; 195-C; 207-C; 216-G; 234-C; 240-G; 321-G; 390-C; 409-A; 612-T

ITS1, 5.8S, ITS-2 gene region with reference to Genbank Accession XXA02: 56-A; 383-T; 839-A; 961-T

### Morphological Description

Body length approximately 3.5 mm, maximum 6 zooids. Body color greyish or slightly tinged pinkish clear. Anterior end abruptly conical, posterior end tail-like with adhesive papillae. Sensory bristles present and ciliary pits present.

Two red stripes present at dorso-anterior end. Amount of pigmentation very bright, almost merging medially to form a dorsal band in some individuals.

Preoral gut present and extending to level of brain. Mouth and pharynx large leading to strait intestine. Nematocysts present throughout the body.

Reproductive system not seen.

### Taxonomic Remarks

Previous research on *Microstomum lineare* has shown the species to be remarkably variable in its morphology. Laboratory experiments have demonstrated that body size and shape differs depending on food availability and reproductive period (Bauchhenss 1971; Heitkamp 1982), the presence or amount of eyespot pigmentation depends on light intensity (Bauchhenss 1971), and even the sexual anatomy changes as the animals develop (e.g. male stylet changes shape, testes change in shape and number; Bauchhenss 1971; Faubel 1974). Similarly, body color has been reported as colorless, white, yellow, grey, reddish or brownish (Luther 1960). The wide ranges of morphological variation combined with a general paucity of morphological features makes defining significant morphological differences between species of the *M. lineare* complex difficult. The ranges of body size, shape and coloration of all species overlapped with each other and with previous descriptions of *M. lineare* (Fig. 4). Similarly, sensory bristles, cilia, ciliary pits, and digestive system with preoral gut were present and consistent across species and with previous records.

We recorded some morphological differences between the new species, e.g. the extent of nematocysts in the body, the number of zooids and the eyespot pigmentation, although it appears that each of the new species was within the normal range for the highly variable *M. lineare* (see, for instance, Fig. S4). The morphological diversity of *M. lineare* implies there may be a similar degree of variation also within the three new species. However, too few individuals were collected of each species to provide an accurate measurement of their morphological variability at this time.

## Discussion

### Cryptic lineages

Our results split *Microstomum lineare* into four divergent clades in all analyses except in the 18S gene phylogeny. However, 18S has been shown to be an inappropriate molecular marker for distinguishing closely related meiofaunal species because it greatly underestimates species diversity even when compared to estimates based on morphology (Tang *et al*. 2012). Taken with results from bPTP tests, our analyses indicate that the four lineages are separate cryptic species.

An additional 18S sequence from a specimen identified as *M. lineare* was downloaded from GenBank, and it differed from the 18S sequences of all other *M. lineare* specimens by approximately 1% (12-15 bp) and may therefore represent an additional cryptic species. This specimen (Genbank Accession D85092) was reportedly collected from Okayama, Japan (Katayama *et al*. 1996). However as no other information on the morphology, ecology or other DNA loci is currently available, we refrain from further formal separation at this time.

### Genetic Diversity

Unsurprisingly, genetic structuring was most evident with the separation of the three specimens collected from Massachusetts, USA (*Microstomum zicklerorum* sp. n.) from the European specimens. The Massachusetts specimens formed a reciprocally monophyletic clade in all ML analyses including the 18S gene phylogeny (Figs. 2, S1-3), and ITS sequences showed F_st_ values at or very near 1.0 between the three specimens and all other *M. lineare* complex groupings (Table S3). The separation of this clade is expected given that large marine waters such as the Atlantic Ocean should constitute an unbridgeable biogeographic barrier for the majority of freshwater free-living flatworms (Artois *et al*. 2011).

However, the remaining three lineages represented in Europe had very little geographic structuring. We found that *M. lineare* collected from very distant locations could nevertheless be genetically similar or identical. Our results showed generally low pairwise F_st_ indicative of high gene flow between locations across Sweden, Finland and Belgium (Table S3), and the majority of specimens collected from such locations formed a single clade with very low gene diversity (Figs. 2, S1-3). Indeed, the most common haplotype (Hap1) occurred in specimens collected from locations as distant as Buresjön in northern Sweden and Hoge Kempen Park in Belgium (Table 1), approximately 1800 km linear distance. The widespread haplotype distributions and lack of geographic structuring contrasts with genetic studies of other zooplankton species (e.g. bryozoans, Nikulina *et al*. 2007; cladocerans, Petrusek *et al*. 2004, Xu *et al*. 2009; rotifers, Fontaneto *et al*. 2008) and strongly suggest that *M. lineare* possesses higher dispersal abilities than previously believed.

*M. lineare* may be particularly well-suited to rapid dispersal due to certain life-history characteristics such as their overall small body size, ability to survive a relatively wide range of environments (Karling 1974) and the production of dormant eggs (Heitkamp 1982). Similarly wide species distributions have been attributed, for instance, to large ranges of salinity tolerances in certain species of freshwater shrimp and mysid crustaceans (Audzijonyte *et al*. 2006; Page & Hughes 2007) or to the desiccation-resistant resting eggs of branchiopods and many other freshwater invertebrates (Havel & Shurin 2004). Further, the ability of *M. lineare* to reproduce asexually through fissioning would greatly aid in colonization of new habitats by avoiding the problem of mate limitation and allowing for clonal propagation (Artois *et al.* 2011).

Larsson *et al*. (2008) measured genetic variation among the catenulid flatworm *Stenostomum leucops* (Platyhelminthes: Catenulida) from localities throughout Sweden and, similar to our results with *M. lineare*, found little genetic variation for 18S, CO1 and ITS sequences. Comparisons between *S. leucops* and *M. lineare* are particularly pertinent as both organisms share many characteristics despite belonging to widely separate Platyhelminthes groups. Both are widespread limnic species common to the same environments and, in fact, are often collected together in the same sample. They are of similar size and body shape (causing them to be easily initially mistaken for each other) and have near identical lifecycles, reproducing through paratomy for the majority of the year followed by short periods of sexual reproduction occurring primarily at the end of the season.

However, unlike our results with *M. lineare*, there were significant genetic distances detected between specimens *of S. leucops* from mainland Sweden and the Swedish island of Gotland, which likely can be explained by differences in the salinity tolerances of the two species*. M. lineare*, being able to inhabit waters with salinity to 8‰ (Karling 1974), occurs commonly in the brackish waters of the Baltic Sea and may be able to freely move between Gotland and the rest of the country. *S. leucops* is a limnic species only or with salinity tolerances to 2‰ (Luther 1960), and the Baltic sea should thus act as a barrier to gene flow. From this, it might also be speculated that while *M. lineare* can cross the Baltic sea to mainland Europe, as demonstrated by the shared Belgium-Scandinavian haplotype, *S. leucops* is likely more restricted.

*Microstomum lineare* is the most common and widespread species of *Microstomum*, which has lead to it being a type of model organism for the genus. The presence of multiple distinct but cryptic lineages in single locations, such as was found in Mustalahti, Finland, Stockholm, Sweden and Hoge Kempen Park in Belgium, implies that more cryptic lineages may be present in other locations as well and may have larger implications for the current knowledge of *Microstomum*. Research on a wide variety of topics, including reproduction (Reuter & Kuusisto 1992; Fairweather & Skuce 1995; Gustafsson et al. 1995), regeneration (e.g. Reuter & Palmberg 1989; Palmberg 1986, 1991; Reuter & Kreshchenko 2004), life cycle and ecology (Bauchhenss 1971; Heitkamp 1982) and nervous and sensory systems (e.g. Palmberg et al 1980; Reuter et al. 1986, 2001; Wilkgren et al. 1986), has been performed on *M. lineare* collected from environmental samples over a period of weeks or, in some cases, years. The presence of multiple cryptic species in a single location means that without genetic testing, there is no guarantee that the collected specimens belonged to the same species, even if they had been collected simultaneously.

### Taxonomy

Traditional taxonomy primarily bases species descriptions on morphological characters and thereby fails to capture the hidden biodiversity in cryptic lineages, yet there is no official requirement by the International Code of Zoological Nomenclature obligating taxonomists to use morphology as the primary data source. The questions of how much data is necessary for taxonomic descriptions and of which type seems to be continually discussed (DeSalle *et al*. 2005; Lipscomb *et al*. 2003; Jörger *et al*. 2012, 2013; Albano *et al*. 2011; Lim *et al*. 2012; Sauer *et al*. 2012; Fontaneto *et al*. 2015), but the value of utilizing DNA data, specifically, to delimit and describe species has been well argued by previous authors (Jörger & Schrödl 2013; Leasi & Norenburg, 2014; Caron, 2013;).

For species such as *Microstomum lineare*, using DNA evidence to demark species is perhaps even more necessary. As noted in the introduction, the animals have relatively few morphological characters on which to base descriptions and laboratory studies have demonstrated very wide amounts of morphological variation within individual lineages. *M. lineare* and species like it lack reproductive organs during periods of asexual reproduction. Moreover, sexual anatomy is known to change as the animals develop (e.g. male stylet changes shape, testes change in shape and number). DNA evidence is arguably more effective than traditional morphological methods in discerning species because it does not depend on environmental conditions or the maturity of the animal.

In order for DNA information to be fully useful to taxonomists, species must be formally described and named, not just delineated, as large numbers of unrecognized candidate species or OTUs leads to confusion and hinders the discovered taxa from being included in future research or conservation efforts (Jörger & Schrödl 2013). This is particularly relevant when the number of metabarcoding-based studies is quickly growing. Formally described species with the relevant sequence data attached can be incorporated into reference databases and enhance the value of metabarcoding surveys. For this reason, we formally describe three new species, *Microstomum artoisi* sp. n., *Microstomum tchaikovskyi* sp. n., and *Microstomum zicklerorum* sp. n. based on our molecular species delineation results and also provide taxonomic sequence data for *Microstomum lineare*.

The diversity within the *Microstomum lineare* species complex revealed by molecular data was not morphologically evident. All specimens had body size, shape and coloration consistent with previous descriptions of *M. lineare* (Fig. 4). While variation in the anterior red eye stripes was found between the different species, amount of eyespot pigmentation has been shown to likewise greatly vary even among individuals of the same lineage depending on light intensity (Bauchhenss 1971; Fig S3). Similarly, all species included specimens with multiple zooids consistent with normal variation in the *M. lineare* species descriptions. Sensory bristles, cilia, ciliary pits, nematocysts, and digestive system (including preoral gut) were present and consistent across species and with previous records. Reproductive organs that were morphologically congruous to previous *M. lineare* descriptions were found in a few specimens only, all from the “main” *M. lineare* complex clade (Fig. S4).

*Microstomum lineare* was originally described by Müller (1773) from an unknown location, and a type specimen was never designated. As Müller was from Copenhagen and much of his work centered on the fauna within and surrounding Denmark, Norway and northern Europe, the clade most widely distributed in this area is the likeliest candidate to include individuals from the original location. The *lineare* clade included specimens from all three of the European countries that were sampled and can be inferred to be widespread throughout northern Europe. We therefore consider the specimens in this clade as *M. lineare*, but we refrained from officially designating a neotype as all specimens were destroyed in the process of DNA sequencing.

## Acknowledgements

There were many individuals who provided help and support while sampling. Many thanks to: Dr. Tom Artois, Edmond Atherton, Adam Bates, Dr. Rick Hochberg, Ylva Jondelius and the entire Zickler family. Thank you also to Dr. Kjell Arne Johanson and Dr. Yngve Brodin for their contributions to identification of gut contents. We are grateful to the staffs of the Stensoffa Research Station, Abisko Scientific Research station, and Umeå Marina Forskningcentrum. This research was supported by a grant from the Swedish Taxonomy Initiative (no. DHA 2014-151 4.3) to UJ.

---

Table S1. Collecting information for the specimens used in this study. Collecting location, collection area abbreviations (see also Figure 1), specific coordinates and dates are given for each specimen, where available. In addition, the GenBank Accession Numbers for 18S, CO1 and ITS (ITS1-5.8S-ITS2) sequences and references are listed.

Table S2: Primer sequences, references and protocols for amplification of 18S, CO1 and ITS1-5.8S-ITS2 sequences.

Table S3: Approximate distances in km between sampling locations (Figure 1, Table S1) and results of the mantel tests, including all collected specimens and specimens from Europe only.

Table S4: Average uncorrected Kimura-2-Parameter (K2P) genetic distances for ITS1-5.8S-ITS2 sequences between (below diagonal) and within (diagonal) locations (See Table S1, Figure 1) and pairwise Fst values (above diagonal). *p<0.05, **p<0.01

Figure S1: 18S gene tree. For clarity, specimens with exactly identical sequences are combined to a single branch. Specimens are listed with their GenBank accession number(s). Bootstrap values are given at each node.

Figure S2: CO1 gene tree. For clarity, specimens with exactly identical sequences are combined to a single branch. Specimens are listed with their GenBank accession number(s). Bootstrap values are given at each node.

Figure S3: ITS1-5.8S-ITS2 gene tree. For clarity, specimens with exactly identical sequences are combined to a single branch. Specimens are listed with their GenBank accession number(s). Bootstrap values are given at each node.

Figure S4: Eye pigmentation variation in *Microstomum lineare*. A. Specimen from Grundträsket, Sweden (GenBank Accession XXXA27) with large amounts of eyespot pigmentation. B. Specimen from Pirttivuopio, Sweden (GenBank Accession XXXA51) with medium amounts of eyespot pigmentation. C. Specimen from Mälaren, Sweden (GenBank Accession XXXE66) with low amounts of eyespot pigmentation. D. Anterior view of specimen from Grundträsket, Sweden (GenBank Accession XXXA27). The animal is “looking up” at the camera so that the typical two dorsolateral eye-stripes and the less typical two ventrolateral eye-stripes are visible. E. Anterior view of specimen from Mälaren, Sweden (GenBank Accession XXXE64). The animal is “looking up” at the camera so that the typical two dorsolateral eye-stripes are visible. Note the absence of ventrolateral pigmentation. F. Eye pigmentation on a developing zooid. Animal collected from Abisko, Sweden (GenBank Accession XXXA63). Arrow points to the mouth opening. Bar measures 100 µm in all images.

Figure S5: Reproductive structures of *Microstomum lineare*. A. Posterior view of male structures. B. Focus on the ovary. C. Focus on the vesicular seminalis and stylet. i intestine; o ovary; st stylet, vs vesicula seminalis. Bar measures 100 µm in all images.

Figure S6: Intestinal contents of *Microstomum lineare*. A. *M. lineare* from Abisko, Sweden (GenBank Accession Number XXXA73) with red diatom in posterior intestine. B *M. lineare* from Mariestad, Sweden (GenBank Accession Number XXXE126) with a chironomid larva in the mid-intestine. C. *M. lineare* from Gevsjön, Sweden (GenBank Accession Number XXXA07) with a chironomid larva in the mid-intestine. D. The same chironomid larva of Fig. S5C after the M*. lineare* ejected it from its mouth. *Microstomum* often eject gut contents from their mouth after being squeezed between a cover slip and a glass slide. E. *M. lineare* from Mariestad, Sweden (GenBank Accession Number XXXE108) with the cladoceran *Chydorus* sp. in its posterior intestine. F. Focus on the same *Chydorus* sp. as in Fig S5E (still in the intestine). G. *M. lineare* from Mälaren, Sweden (GenBank Accession Number XXXE65) with *Chydorus* sp. in its intestine.

