## Supplementary figures and images for "Wide distributions and cryptic diversity within a *Microstomum* (Platyhelminthes) species complex"

### Supplementary Materials

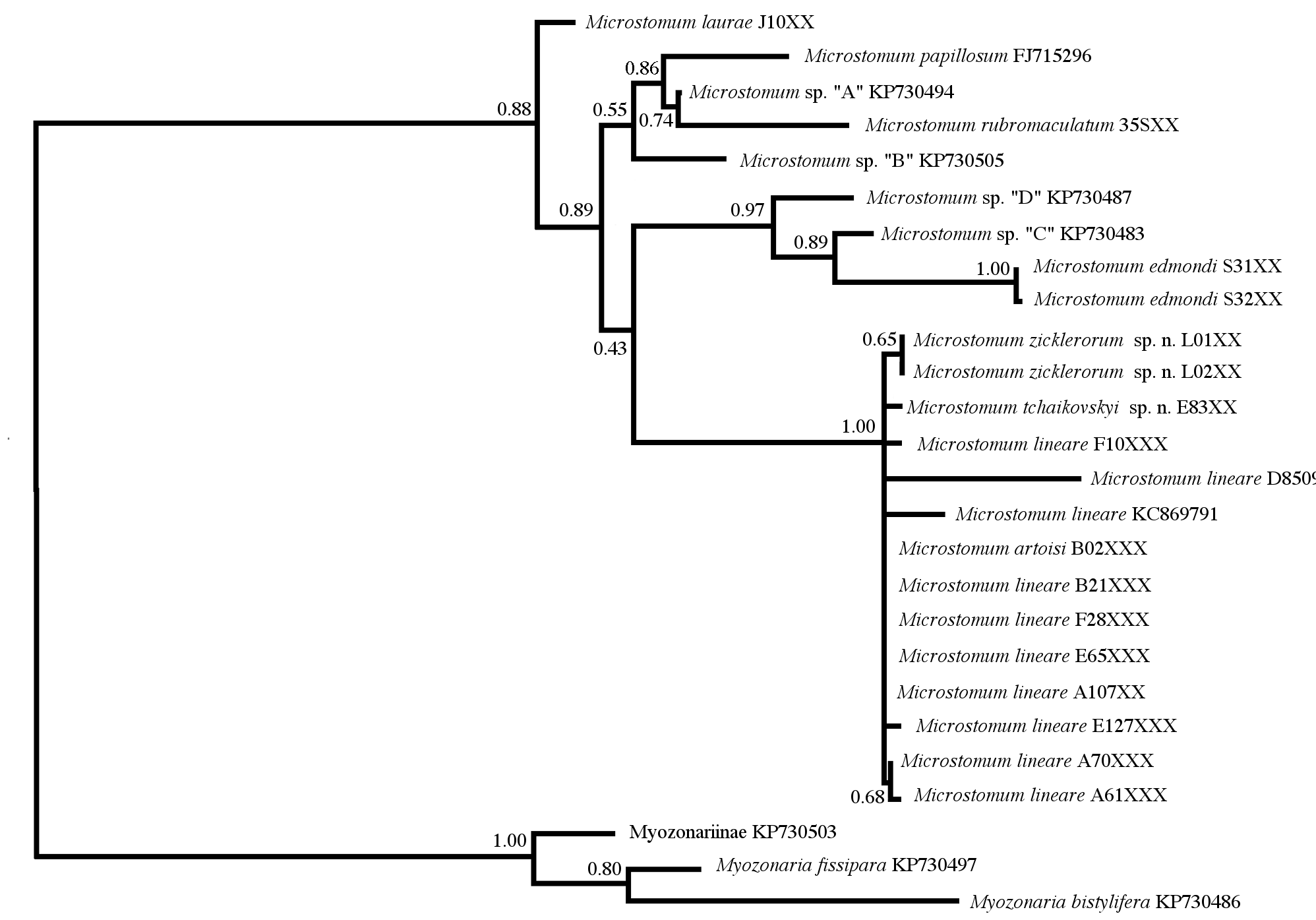

### Supplementary Materials

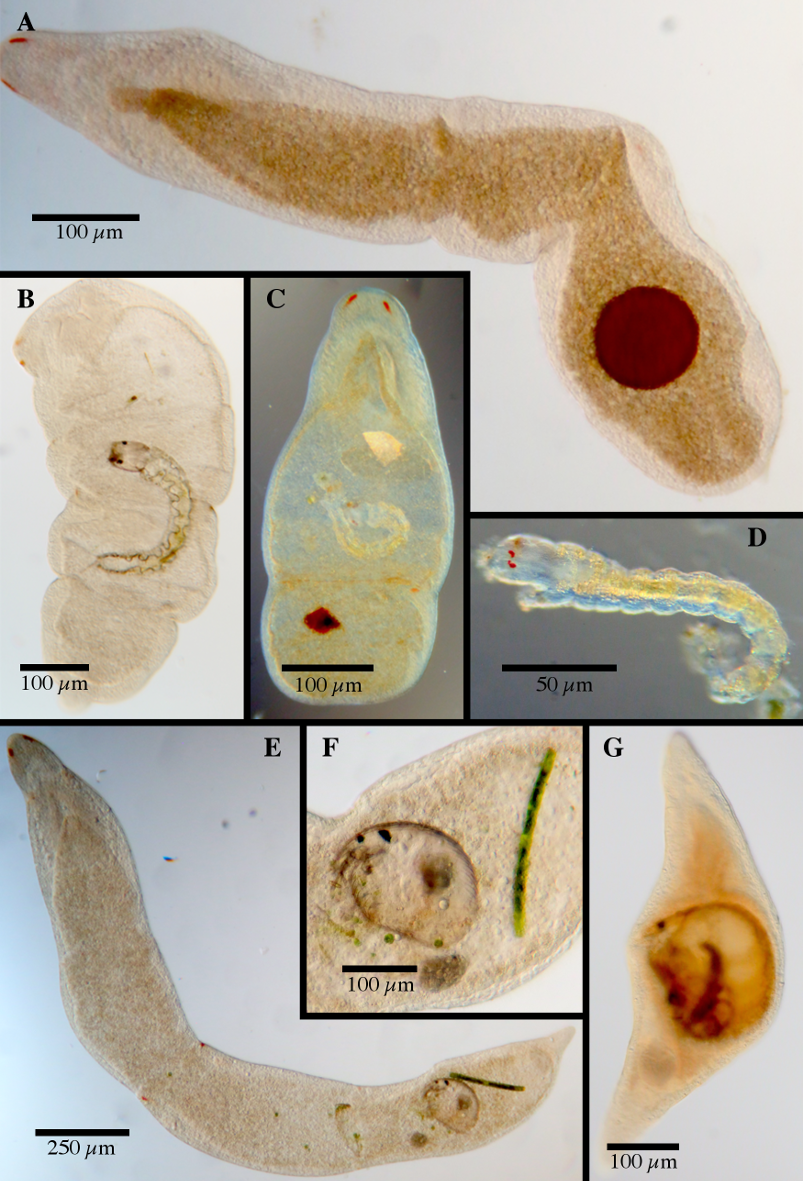

### Supplementary Materials

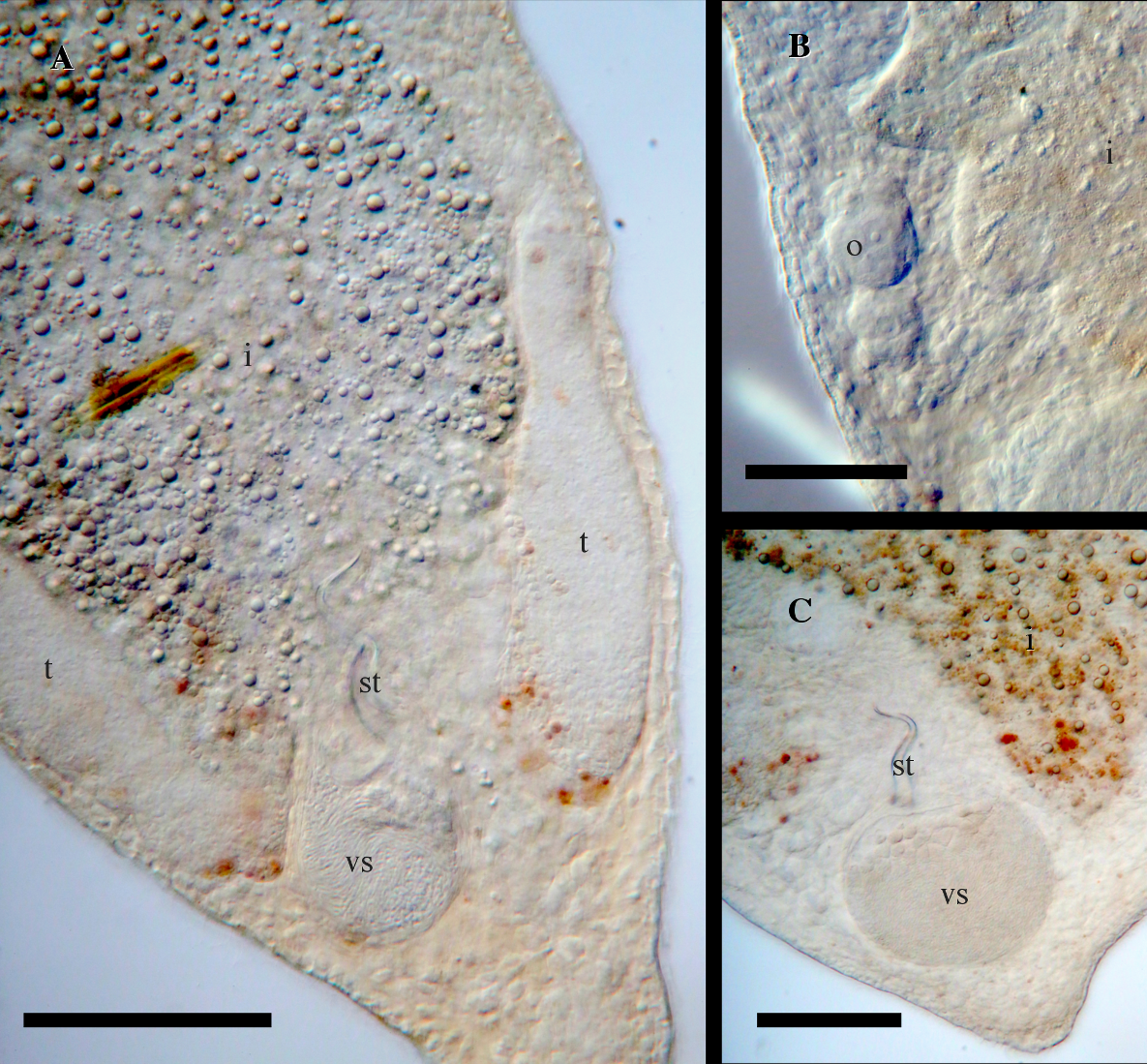

### Supplementary Materials

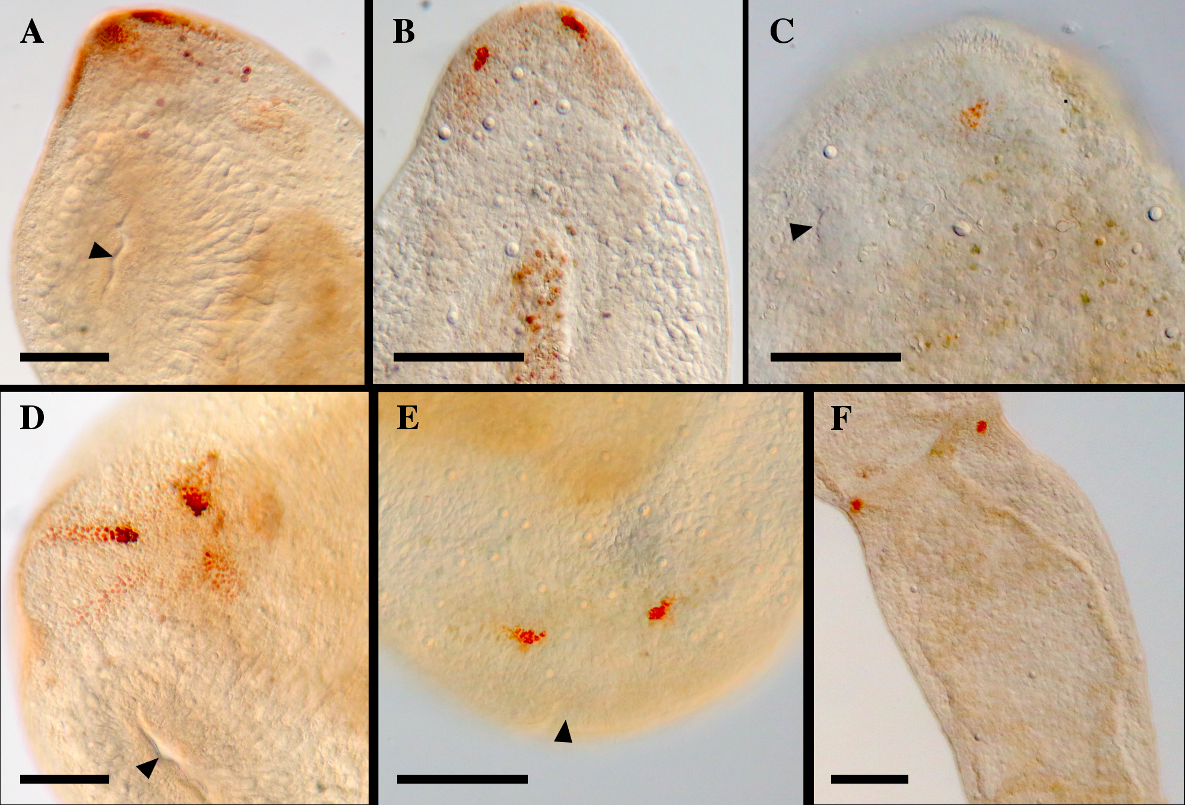

### Supplementary Materials

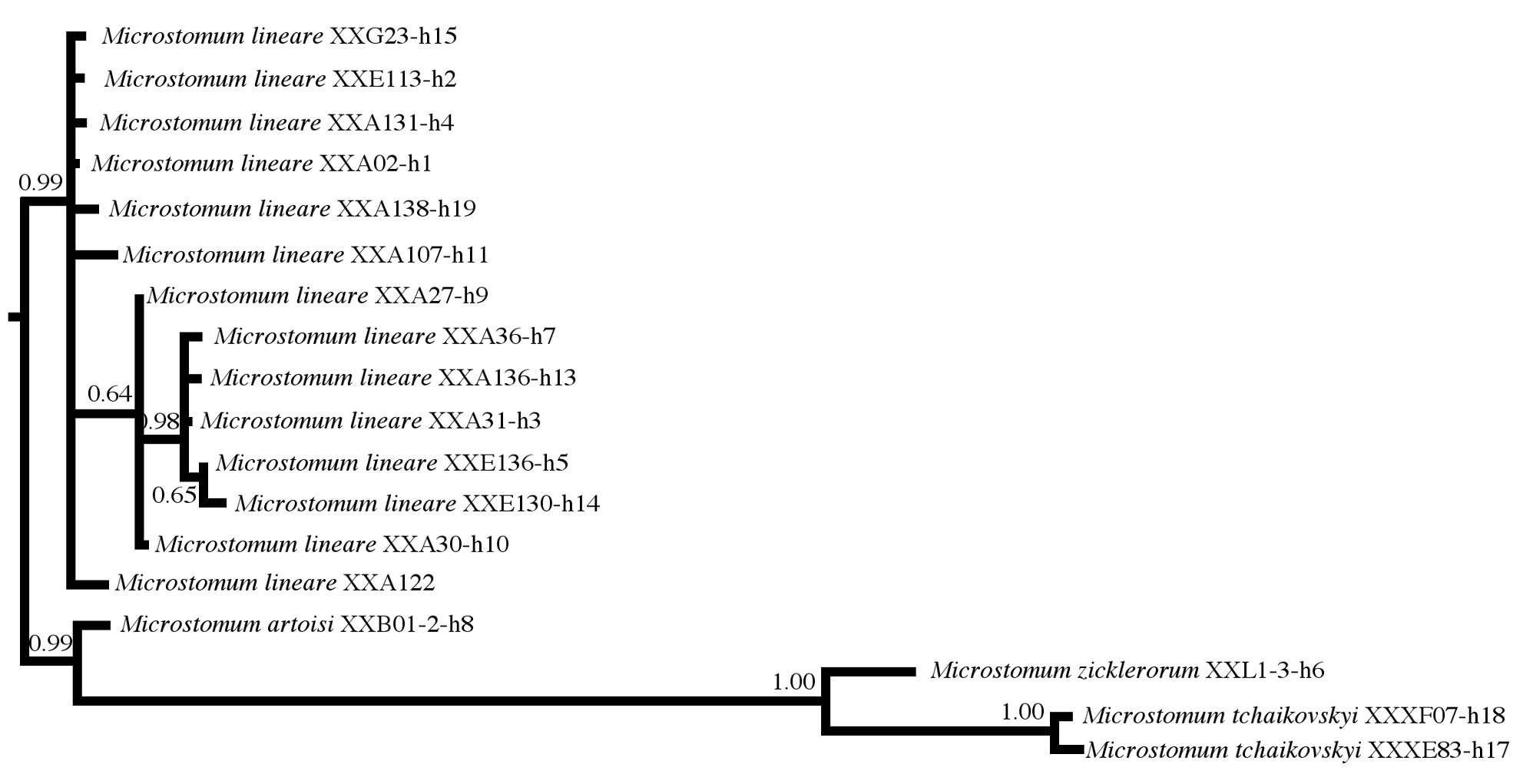

### Supplementary Materials

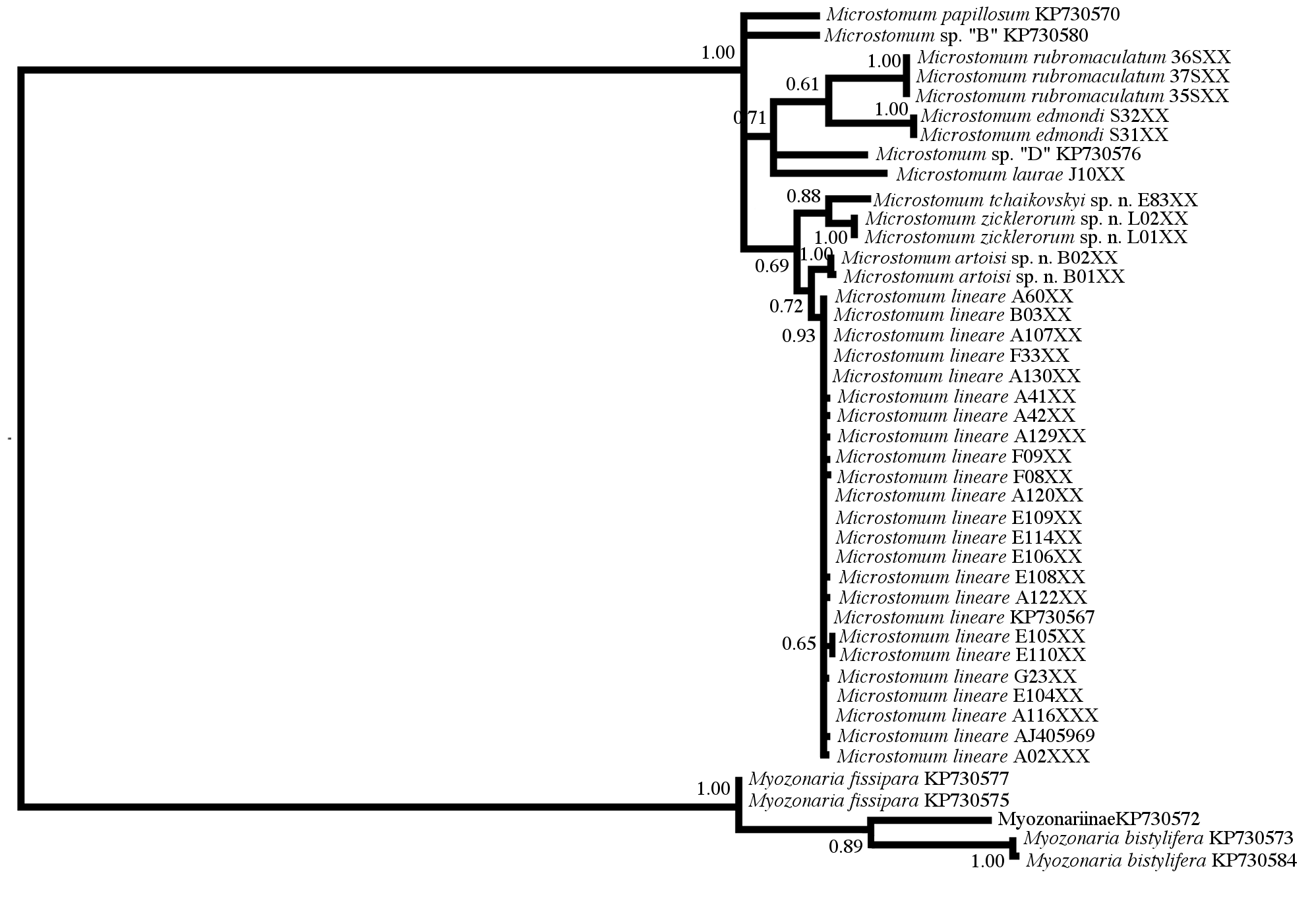
