## Supplementary Materials for "Wide distributions and cryptic diversity within a *Microstomum* (Platyhelminthes) species complex"

Table S2: Primer sequences, references and protocols for amplification of 18S, CO1 and ITS1-5.8S-ITS2 sequences.

Gene

Primer	Direction	Reference	Sequence

Protocol

18S

WormA	Forward	Littlewood & Olsen, 2001	GCGAATGGCTCATTAAATCAG

WormB	Reverse	Littlewood & Olsen, 2001	CTTGTTACGACTTTTACTTCC

1270F	Internal	Littlewood et al. 2000	ACTTAAAGGAATTGACGG

1270R	Internal	Littlewood et al. 2000	CCGTCAATTCCTTTAAGT

5 min at 94°C; 40x (30s at 94°C, 30s at 54°C, 2 min at 72°C); 10 min at 72°C

CO1

Mac_COIF	Forward	Janssen et al. 2015	GTTCTACAAATCATAAGGATATTGG

Mac_COIR	Reverse	Janssen et al. 2015	TAAACYTCWGGGTGACCAAAAAACCA

5 min at 94°C; 5x (30s at 94°C, 90s at 45°C, 60s at 72°C); 35x (30s at 94°C, 90s at 51°C, 60s at 72°C); 10 min at 72°C

ITS1, 5.8S, ITS2

ITS4	Reverse	White et al. 1990	TCCTCCGCTTATTGATATGC

ITS5	Forward	White et al. 1990	GGAAGTAAAAGTCGTAACAAGG

5 min at 94°C; 40x (30s at 94°C, 30s at 56°C, 2 min at 72°C); 10 min at 72°C
