## Supplementary Materials for "Wide distributions and cryptic diversity within a *Microstomum* (Platyhelminthes) species complex"

Table S1. Collecting information for the specimens used in this study. Collecting location, collection area abbreviations (see also Figure 1), specific coordinates and dates are given for each specimen, where available. In addition, the Genbank Accession Numbers for 18S, CO1 and ITS (ITS1-5.8S-ITS2) sequences and references are listed.

| Location | Abbreviation | Coordinates | | Date | Genbank Accession Numbers | | | Reference |
|---|---|---|---|---|---|---|---|---|
| | | | | | 18S | COI | ITS | |
| *Microstomum lineare* | | | | | | | | |
| Tensjön, Sweden | DA | 61°39’36 N | 15°14’52 E | 1-7-2016 | XXX-XX2 | MF185698-9 | XXX-XX2 | Atherton & Jondelius 2017 |
| Gevsjön, Sweden | JÄ | 63°21’42 N | 12°40’40 E | 2-7-2016 | XXX-XX2 | XXX-XX2 | XXX-XX2 | |
| Ängermanälven, Sweden | L4 | 64°26’03 N | 16°47’16 E | 6-7-2016 | XXX-XX1 | XXX-XX1 | XXX-XX1 | |
| Grundträsket, Sweden | L3 | 65°28’29 N | 17°37’45 E | 8-7-2017 | XXX-XX1 | XXX-XX1 | XXX-XX1 | |
| Ahasjön, Sweden | L3 | 65°27’33 N | 17°46’19 E | 8-7-2017 | XXX-XX1 | XXX-XX1 | XXX-XX1 | |
| Norr-svergoträsket, Sweden | L3 | 65°32’21 N | 17°38’26 E | 10-7-2016 | XXX-XX8 | XXX-XX8 | XXX-XX8 | |
| Buresjön, Sweden | L3 | 65°33’24 N | 17°51’44 E | 10-7-2016 | XXX-XX1 | XXX-XX1 | XXX-XX1 | |
| Pirttivuopio, Sweden | L2 | 67°52’11 N | 19°13’07 E | 13-7-2016 | XXX-XX3 | XXX-XX2 | XXX-XX3 | |
| Alavuopio, Sweden | L2 | 67°52’06 N | 19°16’52 E | 13-7-2016 | XXX-XX1 | XXX-XX1 | XXX-XX1 | |
| Laukkujärvi, Sweden | L2 | 67°49’49 N | 19°37’22 E | 13-7-2016 | XXX-XX1 | XXX-XX1 | XXX-XX1 | |
| Pond by E10, Sweden | L1 | 68°20’53 N | 18°58’12 E | 15-7-2016 | XXX-XX1 | XXX-XX1 | XXX-XX1 | |
| Creek by E10, Sweden | L1 | 68°25’41 N | 18°26’44 E | 17-7-2016 | XXX-XX4 | XXX-XX4 | XXX-XX4 | |
| Báktájävri, Sweden | L1 | 68°25’51 N | 18°33’16 E | 17-7-2016 | XXX-XX3 | XXX-XX4 | XXX-XX2 | |
| Stor-Tannörsavan, Sweden | ÅN | 63°26’26 N | 19°39’13 E | 28-7-2016 | XXX-XX3 | XXX-XX3 | XXX-XX3 | |
| Lilla-Tannörsavan, Sweden | ÅN | 63°26’27 N | 19°38’42 E | 28-7-2016 | XXX-XX2 | XXX-XX3 | XXX-XX2 | |
| Bergsjön, Sweden | ÅN | 63°38’27 N | 19°10’37 E | 31-7-2016 | XXX-XX2 | XXX-XX5 | XXX-XX3 | |
| Yttre Lemesjön, Sweden | ÅN | 63°37’53 N | 19°03’37 E | 31-7-2016 | XXX-XX1 | XXX-XX2 | XXX-XX1 | |
| Storsjön, Sweden | ÅN | 63°37’44 N | 18°41’15 E | 31-7-2016 | | XXX-XX1 | XXX-XX1 | |
| Campus Pond, Belgium | B | 50°57’45 N | 05°23’23 E | 20-6-2017 | XXX-XX1 | XXX-XX1 | XXX-XX1 | |
| Hoge Kempen Park, Belgium | B | 51°00’03 N | 05°40’44 E | 20-6-2017 | XXX-XX4 | XXX-XX4 | XXX-XX4 | |
| Valjeviken, Sweden | SK | 56°03’44 N | 14°32’25 E | 5-9-2015 | XXX-XX1 | XXX-XX2 | XXX-XX3 | |
| Mälaren, Sweden | SÖ | 59°20’12 N | 17°53’54 E | 6-10-2015 | XXX-XX4 | XXX-XX6 | XXX-XX3 | |
| Rosjön, Sweden | SÖ | 59°26’26 N | 18°00’03 E | 27-7-2017 | XXX-XX5 | XXX-XX5 | XXX-XX5 | |
| Edsviken, Sweden | SÖ | 59°25’47 N | 17°57’42 E | 30-7-2017 | XXX-XX2 | XXX-XX2 | XXX-XX2 | |
| Mariestadssjön, Sweden | VÄ | 58°42’59 N | 13°50’04 E | 30-7-2017 | XXX-XX9 | XXX-XX22 | XXX-XX21 | |
| Riitalahti, Finland | KE | 62°12’47 N | 24°45’39 E | 5-6-2017 | XXX-XX3 | XXX-XX3 | XXX-XX3 | |
| Mustalahti, Finland | KE | 62°01’31 N | 24°41’03 E | 5-6-2017 | XXX-XX2 | XXX-XX2 | XXX-XX2 | |

| | | | | | | | | |
|---|---|---|---|---|---|---|---|---|
| River in Loimaa, Finland | HA | 60°54’58 N | 23°12’13 E | 5-6-2017 | XXX-XX1 | XXX-XX1 | XXX-XX1 | |
| Haunisten allas, Finland | TU | 60°29’58 N | 22°12’17 E | 10-6-2017 | XXX-XX8 | XXX-XX8 | XXX-XX6 | |
| Tingstädeträsk, Sweden | GO | 57°44’18 N | 18°37’32 E | 9-8-2016 | XXX-XX2 | | | |
| Pond in Lärbro, Sweden | GO | 57°51’03 N | 18°48’52 E | 15-8-2016 | XXX-XX2 | XXX-XX2 | XXX-XX2 | |
| Läppträsket, Finnland | | 60°03’07 N | 23°40’19 E | 6-8-2008 | KP730484 | KP730567 | | Jannsen et al. 2015 |
| Connecticut, USA | | | | 1-9-2009 | KC869791 | | | Laumer & Giribet 2014 |
| Ushimado, Okayama, Japan | | | | | D85092 | | | Katayama et al. 1996 |
| United Kingdom | | | | | | AJ405979 | | Telford et al. 2000 |
| *Microstomum artoisi* sp. nov. | | | | | | | | |
| Hoge Kempen Park, Belgium | B | 51°00’03 N | 05°40’44 E | 20-6-2017 | XXX-XX2 | XXX-XX2 | XXX-XX2 | |
| Campus Pond, Belgium | B | 50°57’45 N | 05°23’23 E | 20-6-2017 | XXX-XX2 | XXX-XX2 | XXX-XX2 | |
| *Microstomum tchaikovskyi* sp. nov. | | | | | | | | |
| Rosjön, Sweden | SÖ | 59°26’26 N | 18°00’03 E | 15-9-2016 | XXX-E83 | XXX-E83 | XXX-E83 | |
| Mustalahti, Finland | KE | 62°01’31 N | 24°41’03 E | 5-6-2017 | XXX-F07 | XXX-F07 | XXX-F07 | |
| *Microstomum zicklerorum* sp. nov. | | | | | | | | |
| Massachusetts, USA | MA | 42°38’13 E | 71°23’31 W | 11-04-2017 | XXX-XX3 | XXX-XX3 | XXX-XX3 | |
| *Microstomum laurae* | | | | | | | | |
| Saltö, Sweden | | 58°52’41 N | 11°06’56 E | 19-6-2016 | XXX-XX1 | MF185712-13 | | Atherton & Jondelius 2017 |
| Saltö, Sweden | | 58°52’29 N | 11°08’41 E | 17-6-2016 | XXX-XX1 | | | Atherton & Jondelius 2017 |
| *Microstomum rubromaculatum* | | | | | | | | |
| Fiskebäckskil, Sweden | | 58°14’59 N | 11°26’45 E | 20-8-2015 | | XXX-XX2 | MF185693-5 | Atherton & Jondelius 2017 |
| *Microstomum edmondi* | | | | | | | | |
| Munkedal, Sweden | | 58°27’31 N | 11°41’10 E | 15-8-2015 | XXX-XX2 | MF185701-02 | | Atherton & Jondelius 2017 |
| *Microstomum papillosum* | | | | | | | | |
| Koenighafen, Germany | | 55°02’24 N | 08°23’52 E | 8-3-2007 | FJ715296 | KP730570 | | Scharer et al. 2011; Jannsen et al. 2015 |
| *Microstomum* sp A | | | | | | | | |
| Fetovaia Bay, Italy | | 42°43’36 N | 10°09’33 E | 26-4-2010 | KP730494 | | | Jannsen et al. 2015 |
| *Microstomum* sp B | | | | | | | | |

| | | | | | | |
|---|---|---|---|---|---|---|
| Mangrove Bay, Egypt | 25°52’15 N | 34°25’04 E | 11-1-2009 | KP730505 | KP730580 | Jannsen et al. 2015 |
| *Microstomum* sp C | | | | | | |
| Sant Andrea Bay, Italy | 42°48’31 N | 10°08’30 E | 26-4-2010 | KP730483, 93 | | Jannsen et al. 2015 |
| *Microstomum* sp D | | | | | | |
| Pianosa, Italy | 42°34’29 N | 10°03’59 E | 30-4-2010 | KP730487 | KP730576 | Jannsen et al. 2015 |
| *Myozonaria fissipara* | | | | | | |
| Sant Andrea Bay, Italy | 42°48’31 N | 10°08’30 E | 30-4-2010 | KP730497-8 | KP730575, 77 | Jannsen et al. 2015 |
| *Myozonaria bistylifera* | | | | | | |
| Fetovaia Bay, Italy | 42°43’36 N | 10°09’33 E | 26-4-2010 | KP730486 | KP730573 | Jannsen et al. 2015 |
| Sant Andrea Bay, Italy | 42°48’31 N | 10°08’33 E | 26-4-2010 | KP730510 | KP730584 | Jannsen et al. 2015 |
| Myozonariinae | | | | | | |
| Pianosa, Italy | 42°34’29 N | 10°03’59 E | 30-4-2010 | KP730503 | KP730572 | Jannsen et al. 2015 |
